# Pilifers provide rapid mechanosensory feedback during floral tracking in hawkmoths

**DOI:** 10.64898/2026.10.11.758322

**Authors:** Meghna Anand, Mauro Torres, Bingni W. Brunton, Thomas L. Daniel, Alison I. Weber, Tanvi Deora

## Abstract

Hawkmoths expertly hover over flowers during feeding, accurately tracking floral movements even as flowers gently sway in the wind. They use visual and mechanosensory feedback to adjust their position, while their long, flexible mouthpart, the proboscis, stays inserted inside the nectar spur. Although much is known about how hawkmoths detect and process visual motion, far less is understood about how hawk-moths detect mechanical motion cues. Pilifers are hair bundles at the proboscis-head joint, well-positioned to provide feedback about floral motion. Here, we tested if pilifers could provide mechanosensory feedback essential for successful floral tracking. We computed bending at the proboscis base during floral tracking in freely flying hawkmoths and found that floral movements induce small yet significantly correlated lateral bending. Recordings from pilifer afferents showed that they respond to lateral motions of amplitudes similar to those seen in natural floral tracking behaviors. They respond to different directions and speeds of proboscis motion with an average latency of 1.9 ms. Neural tracing revealed extensive sensory arborizations in the sub-esophageal ganglion of the brain and ventral nerve cord, indicating a feedback pathway to the neck and flight motor system that enables rapid coordination of appendages and body for successful feeding.

## 1 Introduction

Flying insects use multisensory feedback to precisely control their movements. This feedback is particularly important in the complex process of nectar feeding by hovering hawkmoths. Once hawkmoths are close to a flower, they guide their long, flexible proboscis along visual, tactile, and chemical cues on the floral surface and target the tiny opening that leads to the nectar reward [1–7]. With their proboscis inserted into the nectar spur, they successfully hover in front of the flower even as the flowers sway in the wind [8]. Both visual and mechanosensory motion cues from the swaying flower guide their body movements, enabling successful flower tracking [9]. These goal-directed movements require precise information about the position of the moth relative to the moving flower.

Multiple types of mechanosensors on the insect body provide proprioceptive and exteroceptive feedback [10–12]. These sensors include: 1) dome-shaped strain sensors called campaniform sensilla on the surface of the exoskeleton, 2) hair bristle plates extending between joints, and 3) chordotonal organs that are located internally in the joints of appendages. These mechanosensory structures respond to strains in the cuticle, deflection of bristles, or stretching of joints. Movement of appendages during locomotion activates these sensors and hence provide feedback that can be used to control locomotion and maintain posture [13–15].

One potential mechanosensory organ well placed to detect relative motion during floral tracking is a pair of hair bristle bundles at the proboscis-head joint, called the pilifers (Figure 1A). They are found on a slight protrusion of the head, dorso-lateral to the proboscis on either side. Based on their position and appearance, several studies have suggested that pilifers could provide proprioceptive feedback during flower handling [9, 16, 17]. Interestingly, in some species of hawkmoths, such as *Acherontia atropos*, pilifers respond to ultrasonic sounds within the range of bat’s echolocation calls, leading to another hypothesis that pilifers are auditory sense organs [18]. A single chordotonal sensory cell at the base of the pilifer of the hawkmoths of the tribes *Macroglossini* and *Acherontiini* is the likely sensory structure responsible for the auditory sensitivity [19, 20]. However, pilifers of some hawkmoth species, such as *Panogena lingens*, have hair bristles and chordotonal sensory cells, but they do not respond to ultrasonic vocalizations [20]. Although these studies highlight the auditory capabilities of pilifers, the hair bristles of pilifers might also be sensitive to other mechanosensory inputs, such as bending of the proboscis during floral probing and tracking, which would directly deflect the bristles and hence provide feedback about floral motion.

**Figure 1.**
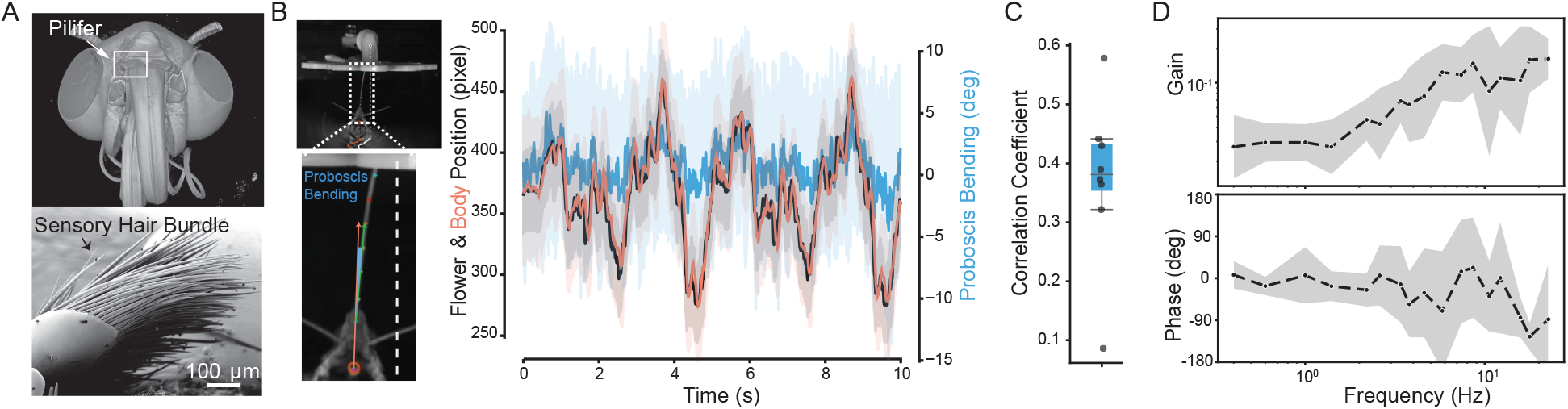
Bending at the proboscis base is informative about flower movements. (A) A micro-computed tomography scan of the hawkmoth head, highlighting the location of the pilifers at the head and proboscis joint. The lower scanning electron microscope image shows a zoomed-in view of individual hair bristles that make up the mechanosensory pilifer bundle. (B) Raw trajectory of hawkmoth body (orange) as it tracks the moving artificial flower (black) resulting in bending at the proboscis base (blue). The solid line and shaded regions represent the mean and std respectively across 8 moths. We computed proboscis bending as the angle between the body heading vector (orange) and proboscis vector (green). Inset is a still from the video showing the proboscis tracked along its length (colored markers). (C) Correlation coefficient between the floral movements and bending at the proboscis base. Each dot represents a 10-second flight bout for an individual hawkmoth (N = 8 hawkmoths). (D) Bode plot showing the gain and phase of the proboscis bending with respect to flower movements.

Although both visual and mechanosensory feedback alone are sufficient to drive successful floral tracking, hawkmoths weigh mechanosensory feedback 2–3 times more than visual feedback [9]. Despite its importance, the mechanosensory organ that provides this high gain and low latency feedback remains unknown. In this study, we investigated whether pilifers are sensitive to lateral movements of the proboscis, such as those that occur during flower tracking. Using high speed videography, we quantified the lateral bending of the proboscis at the head-proboscis joint in freely flying nocturnal hawkmoths that successfully hover while feeding from moving artificial flowers. We find that although the proboscis bending is small, it is significantly correlated with the flower movements relative to the hovering moth. Using extracellular electrophysiological recordings, we show that pilifer afferents respond to proboscis motion within the range of 5–10 degrees, with average latency of a couple of milliseconds. Tracing pilifer arborizations into the brain reveals extensive projections into the ipsilateral sub-esophageal zone (SEZ), an area in the ventral brain responsible for sensorimotor integration of the mouthparts and legs, as well as a few projections into the contralateral SEZ and into the neck connective, which carries feedback to the flight motor system. Taken together, these results suggest that pilifers could play a role in mediating rapid coordination and corrective flight control during flower tracking.

## 2 Methods

### 2.1 Behavioral analysis of proboscis bending during floral tracking

We used previously published videos of hawkmoths tracking artificial flowers to quantify proboscis bending during natural behaviors [9]. We tracked a total of 7 keypoints using a trained DeepLabCut model (version 2.2.2, [21]): 5 along the proboscis length; from the base to where it inserts into the nectary, one on head-proboscis joint, and one on the artificial flower. We fit a cubic polynomial to the proboscis keypoints and computed the tangent to this curve at the base. The angle between the tangent and the heading angle was defined as the proboscis bending angle (Figure 1B; for details see supplementary methods). We computed the cross correlation between the flower’s lateral motion and proboscis bending. In addition, we computed the gain and phase between different input-output pairs of timeseries trajectories, such as the flower motion to proboscis bending, proboscis bending to body motion, and flower motion to body motion. We also quantified the mutual information between flower and proboscis bending across different frequencies. All analyses were performed using custom written code in Python found at https://github.com/TanviDeora/PiliferBehavior.

### 2.2 Electrophysiological recordings from pilifer afferents

We recorded from 15, 2–3 day-old adult *Manduca sexta* from the University of Washington’s moth colony. The moths were cold anesthetized at 4°C and restrained in a custom designed holder. The labial palps were cut to expose the pilifers and a 16-channel silicon probe (Cambridge Neurotech P-1) was inserted at the base of the left pilifer. The silicon probe was connected to a headstage and amplifier, and neural signals were digitized at 30 kS/s (Intan Technology, RHD2132 headstage and RHD2000 recording system). The proboscis was cut near the knee bend (a region where the proboscis naturally bends when uncoiled) and attached to the stepper motor via a custom made adapter and driven by a custom built PID controller. The proboscis was driven with a 10 second long band-limited Gaussian white noise (1–300Hz, at amplitudes ranging from 1–3V), repeated 30 times, with a 1.1 second gap before and after each stimulus repeat. Neural activity from one or more channels from 6 animals was spike sorted into 16 individual units using Plexon Offline Sorter software. Data were analyzed in custom written codes in Python found at https://github.com/MeghnaAnand5/PiliferEphysAnalysis. We used a 20 ms stimulus preceding each firing event to compute the average stimulus feature, the spike triggered average (STA) and nonlinear decision function (NLD) ([22, 23] for details, see supplementary methods). Out of these 16 units, we selected 12 units for further analysis based on the following criteria: a) they should have a sufficient number (*>*250) of firing events, b) the amplitude of the spike triggered average (STA) should exceed one standard deviation of prior triggered ensemble. From these fits, we computed the latency of response as the time delay between the firing event and when the STA first crosses the prior ensemble’s 1 std.

### 2.3 Neural tracing

We used 1–3 day-old adult *Hippotion sp*. and *Daphnis nerii* caught from wild or raised in the colony at Shiv Nadar University, Delhi-NCR. We used closely related nocturnal species within the family Sphingidae for neural tracing due to the unavailability of *M. sexta* in India. We expect their pilifer projections to be similar because these moths are closely related and share similar feeding ecology (nocturnal, hover while feeding from flowers). We cold anesthetized and restrained the hawkmoths in a custom designed holder and exposed the pilifers by cutting the labial palps. We shaved the hair bristles and applied concentrated Texas Red tagged dextran dye (Invitrogen, D3328). The preparation was incubated for 24–72 hrs at RT, fixed in 4% paraformaldehyde solution at 4°C overnight and dissected to expose the nervous system. The nervous tissue was serially dehydrated, cleared and mounted in methylsalicylate, and imaged using confocal microscope (Nikon Ti2E, AXR NSPARC, see supplementary methods for details).

## 3 Results

### Bending at the proboscis base is informative about floral movements

As a hawkmoth feeds from a moving flower, its proboscis is pulled along by the swaying flower. Any mismatch between the position of the hovering moth and the flower results in bending of the proboscis. Since the proboscis is a long, unjointed appendage, movement of the tip will induce bending along the entire length, including at its base, where the pilifers are located. However, the proboscis is highly flexible, and the bending at the base might not necessarily be informative of the movements at the tip. To understand if bending at the proboscis base is proportional to the flower movement, we computed the magnitude of bending in freely flying hawkmoths as they track swaying artificial flowers (videos taken from previously published data [9]). We tracked the position of the artificial moving flower and the moth’s body (Figure 1B, black and orange) as well as the bending of the proboscis near the proboscis-head joint (Figure 1B, blue).

Hawkmoths continuously track the large lateral movements of flowers with high fidelity and minor delays ([9], Figure 1B, Supplementary Figure 3). As a result, the bending at the proboscis base is small; on average, the maximum amplitude of bending is 20.06 *±* 6.02 degrees (mean *±* std, n = 8 moths). Each individual moth approaches the flower at a different angle, which is maintained during the flight bout (Supplementary Figure 2). Hence, even though hawkmoths align their hovering position with the moving flower nearly perfectly, we see variability in the resulting bending at the proboscis base across animals, which is largely dependent on approach angle. Despite the small amplitude of bending, the average correlation coefficient between flower movement and proboscis bending is 0.37 *±* 0.14 (mean *±* std for the Pearson correlation coefficient, *p <* 0.01 for all moths when testing for non-correlation, N = 8 moths, Figure 1C), indicating a significant positive correlation between flower movement and proboscis bending. The artificial flower is actuated at distinct frequencies, each of which is reflected in the deflections at the proboscis base (Supplementary Figure 1). Higher gains between flower motion and proboscis bending at higher frequencies indicate that the higher frequency movements are better represented in the pilifer bending at the head-proboscis joint (Figure 1D and Supplementary Figure 3 and 4). Hawkmoths do not track flowers moving at frequencies beyond 20Hz well, so we could not test how proboscis mechanics shape pilifer inputs at frequencies higher than 20Hz. However, it is important to note that the proboscis possibly bends and vibrates at higher frequencies during other tasks, such as tracking textural gradients on floral surfaces during nectary localization.

**Figure 2.**
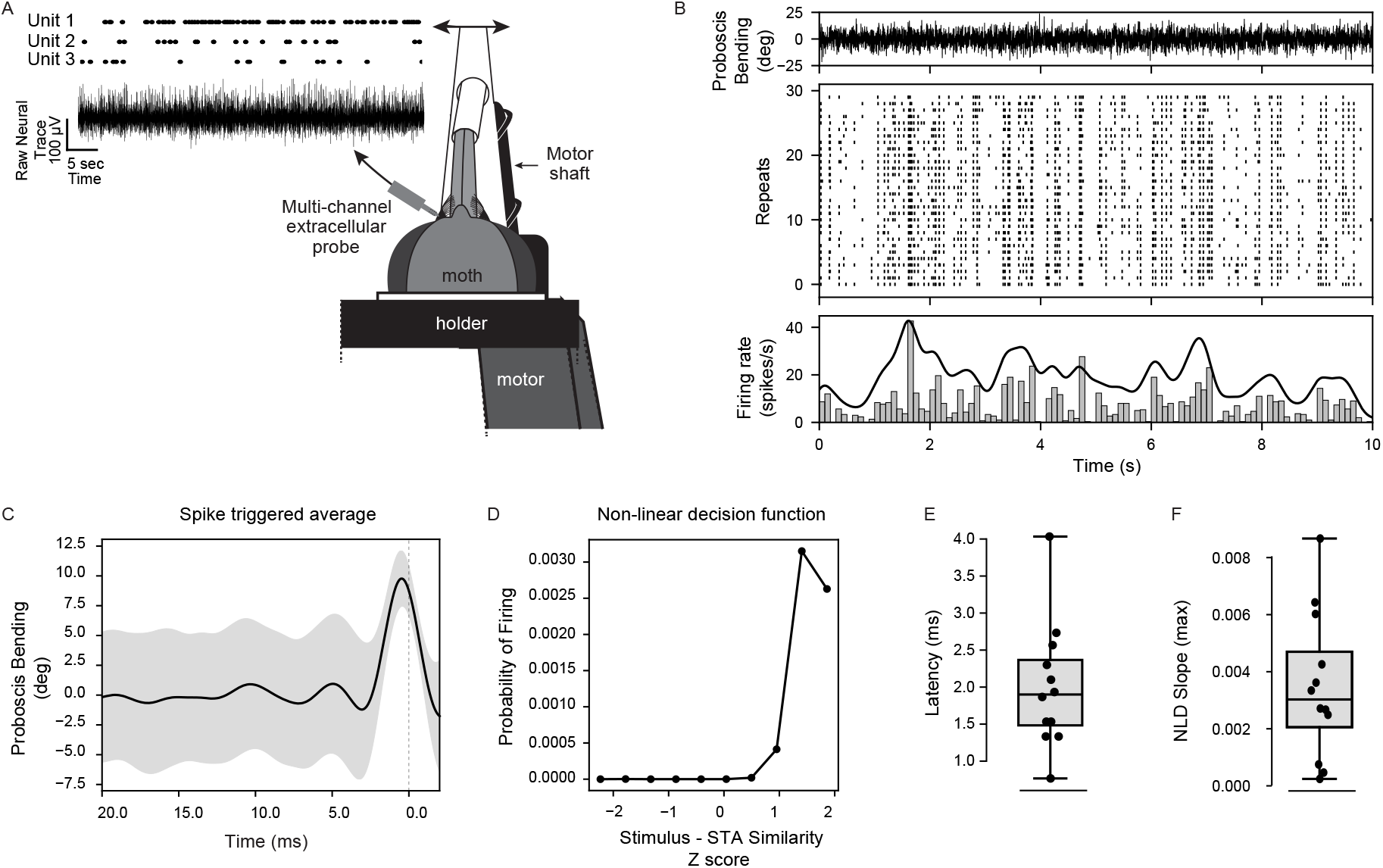
Pilifer neurons respond to lateral motion of the proboscis. (A) An illustration of the electrophysiology setup used for recording from a restrained hawkmoth using a multisite extracellular probe. A representative raw extracellular recording from pilifer nerve afferents is shown in black, which is spike sorted into individual units. (B) Response of one representative unit. *Top:* Proboscis bending at its base. *Middle:* Raster showing the timestamps for all spiking events during the 10 seconds stimulus, across 30 repeats. *Bottom:* Peri-stimulus time histogram showing the average spiking across all repeats. (C) The mean stimulus feature (spike triggered average, STA, in black and the standard deviation in shaded gray) and (D) non-linear decision function (NLD) for the same representative unit shown in B. (E) Response latency and (F) the maximum steepness of the NLD function across all units (N = 12).

**Figure 3.**
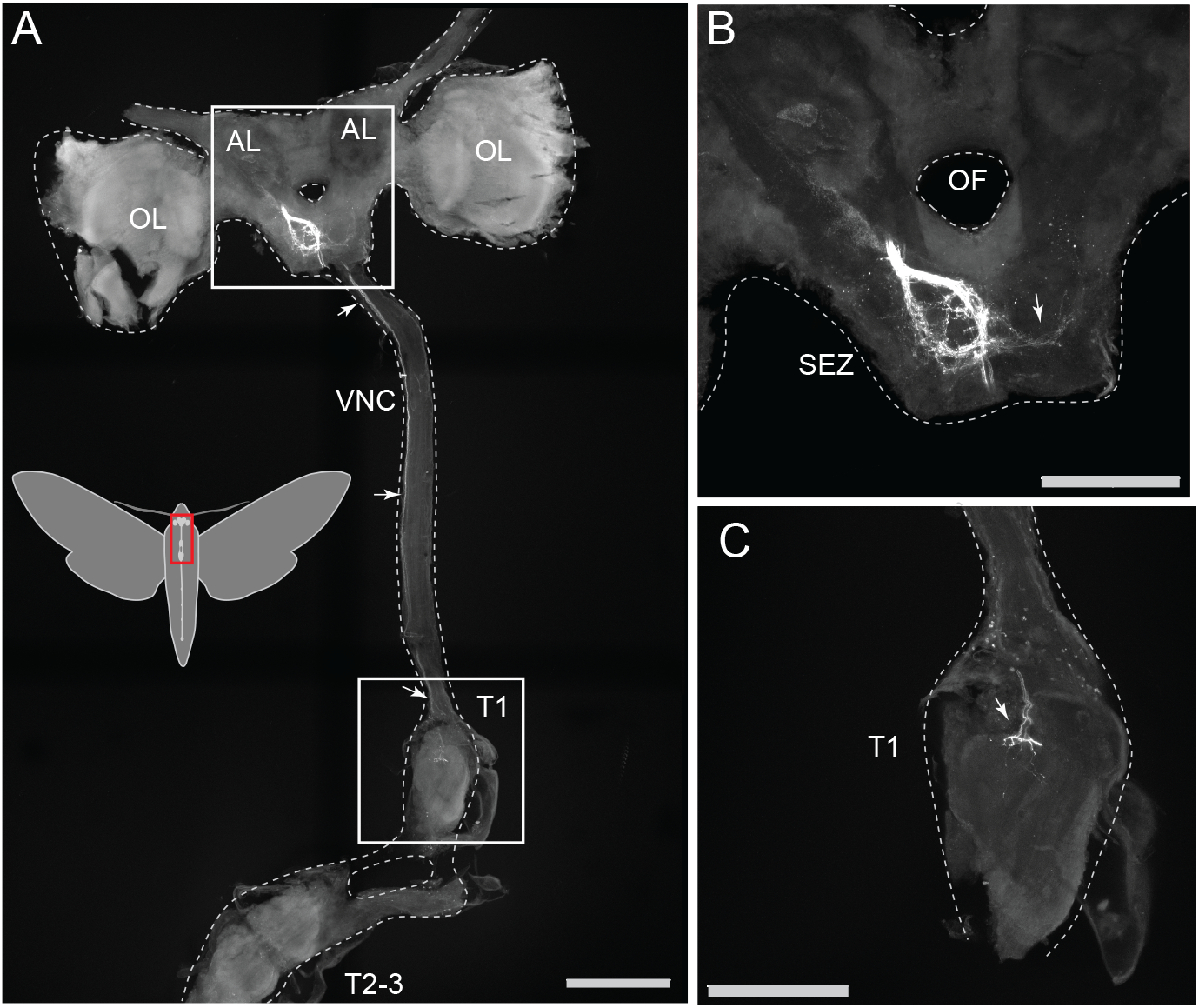
Axonal projections of pilifer hair bristles into the nervous system. (A) Imaging of the brain and thoracic ganglion (highlighted in red on the inset) shows pilifer afferents of *Daphnis nerii* enter the brain and arborize heavily in the ipsilateral sub-esophageal zone (SEZ), with some projections into the contralateral side. (B) shows the zoomed-in view of the SEZ, and (C) shows arborization in the first thoracic ganglion. OL - optic lobe, AL - antennal lobe, VNC - ventral nerve cord, T1 - first thoracic ganglion, T2-3 - second and third thoracic ganglion, OF - oesophageal foramen, SEZ - sub-esophegeal zone. The scale bar represents 500 microns in A and 250 microns in B and C.

### Pilifers respond to lateral movements of the proboscis

We recorded from pilifer afferents to test if pilifers are sensitive to lateral movements of the proboscis. We recorded the response of pilifers in restrained moths to a 10 second band-limited white noise motion stimulus with a multichannel extracellular silicon probe, and spike sorted neural activity from distinct units (16 units from 6 moths, Figure 2A). All recorded units increased activity during the motion stimulus. In response to repeated presentation of the same motion stimulus, individual units exhibited consistent increases in firing rate at specific time points, suggesting that they respond to specific features of the stimulus (Figure 2B).

We performed a reverse correlation analysis to quantify the specific stimulus feature to which each neuron responds (Figure 2C & D, [22, 23]). Of the 16 identified units, 12 showed a distinct feature in the spike triggered average (STA) whose peak deviated significantly from the average random stimulus (see Supplementary Figure 5 for more units). For 9 units, we could compute the range of proboscis deflection during the stimulus, and on average, the units responded to deflections of 9.6 *±* 2.2 degrees (mean *±* std). Units within the left pilifer were sensitive to either leftward or rightward movement. The typical response latency, calculated as the time interval between the firing event and when the STA crosses 1 std of the prior distribution, was 1.9 ms (median; 25th percentile: 1.48 ms and 75th: 2.37 ms, n = 12 units, Figure 2E). The frequency of these stimulus features was between 50–300 Hz, suggesting a broad tuning across the tested range (Supplementary Figure 6). In addition to the STA, we also computed the non-linear decision function (NLD), which reflects a unit’s sensitivity to the average stimulus feature. The representative NLD in Figure 2 shows a sharp increase in the probability of firing as the similarity score between stimulus and STA increases, suggesting that this unit is highly selective. The maximum steepness of the NLD function was typically 0.003 (median) but showed large variation around this value, suggesting that some units are highly selective for the STA feature (high slope) while others are relatively less selective (low slope; Figure 2F).

### Neural tracing reveals extensive arbors in the sub-esophageal zone and ventral nerve cord

We traced the axonal projections of pilifer afferents by back-filling the sensory neurons with fluorescent tracers (Figure 3). We find that the axons of pilifer hair bristles arborize extensively in the ipsilateral sub-esophageal zone (SEZ), with a few contralateral projections. Axons travel along the characteristic circular neural tracks in the SEZ before entering the contralateral side. A subset of the projections also extend into the ventral nerve cord and arborize into the first thoracic ganglion (T1). Although we did not observe projections further than T1 in our neural fills, it is possible that the axonal terminals of these neurons project further down to the second thoracic ganglion as well but were not observed due to limited time of dye diffusion and concentration. Direct projections into the SEZ and tracks into the ventral nerve cord suggest direct, rapid communication to motor neurons of the neck in the SEZ and motor neurons for the neck and wings in the thoracic ganglia.

## 4 Discussion

In this study, we asked if pilifer could provide mechanosensory motion feedback during floral tracking. We show that small bending induced at the proboscis base during flower tracking is significantly correlated with flower movements (Figure 1). This bending is large enough to deflect the hair bristles, which are sensitive to small amplitude motions of the proboscis (Figure 2). Finally, the rapid response latency of pilifer neurons and their arborization in the brain and the thoracic ganglion support their likely role in control of hovering flight during feeding (Figure 3).

Neural recordings demonstrate that units in the left pilifer are sensitive to rapid deflections in both leftward and rightward directions relative to a mean position aligned with the body axis (Supplementary Figure 5). Interestingly, behavioral analysis reveals that moths typically do not keep their body centered with respect to the flower during tracking (Supplementary Figure 2). Rather, moths approach flowers at a random angle and maintain this angle consistently throughout the flight bout, resulting in a mean offset in the proboscis deflection over the course of flower tracking (Supplementary Figure 2, mean angle of 5.1 degrees, N = 8 animals). Future studies will reveal if pilifer mechanosensors encode absolute deviation of the proboscis or if their responses are independent of the baseline angle of the proboscis with respect to the body, or if different units show mixed sensitivity to relative versus absolute position.

We examined neural responses to 5–20°deflections, relevant to floral tracking. We found that neurons were sensitive to deflections on the order of 5–10°. However, they may also detect smaller-amplitude vibrations during floral exploration, potentially at different frequencies than those tested. Hawkmoths use tactile cues such as the shape and texture of floral surfaces to localize and target the nectary opening [1, 3]. Proboscis tip’s interaction with floral surfaces will likely induce bending and vibrations along its entire length. Although hair bristle STAs suggest sensitivity to 5–10° deflections, the associated chordotonal neuron may detect smaller vibrations during floral exploration and nectary targeting.

Hair bristle sensors occur at joints, such as the legs and antennal base, and across other body regions, including wing edges, haltere tips, tegula, and head. [11, 24–27]. Hair bristles occur in regular arrays within a hair plate, such as on insect legs and hawkmoth antennae, or singly/pairs on structures such as halteres and wings. The pilifer bristles form a dense cluster, all leaning towards the proboscis, with length increasing medially to laterally. We find neurons responding to leftward or rightward movement within the same hair bundle, although the basis of this direction tuning is unclear. In locusts’ cephalic wind sensors, directional tuning is determined by the shape of the cuticular collar, and curvature of the hair [27], while cricket cercal sensilla’s tuning to motion deriatives is dependent on hair length; shorter hair encode changes in acceleration and longer hair encode changes in velocity [28, 29]. Pilifer bristles may similarly differ in sensitivity to position, velocity, acceleration, or motion frequency. Although data in Supplementary Figure 6 supports this, further data are needed to access population-level frequency tuning.

Pilifer neurons project into several regions of the nervous system, with the densest projection into the ipsilateral SEZ, and sparser projections into the contralateral SEZ and the ventral nerve cord. This spatially distinct projection pattern may indicate functional specialization among pilifer neurons. In mechanosensory systems such as the chordotonal organs (CO) in fly leg, distinct sensor subsets encode different movement features and project to different regions of the ventral nerve cord, suggesting functionally segregated channels for processing different movement features [30]. Subsets of wing and haltere strain sensors similarly project to different areas within the thoracic ganglion [31, 32]. Pilifer bristles may therefore also be functionally specialized, with different subsets supporting distinct behaviors.

In summary, hawkmoths rely on multimodal sensory feedback from vision, mechanosensation, and olfaction to hover and feed from flowers. Our results identify pilifers as the mechanosensor involved in floral tracking. The rapid response of pilifer mechanosensors could provide the fast feedback required for flight control, complementing the slower visual cues. These results raise interesting future areas of investigation such as how bristle mechanics shape sensory encoding, and how pilifer feedback is integrated with other modalities to coordinate proboscis proboscis and body movements and floral tracking. These findings highlight the importance of mechanosensory feedback in hawkmoth feeding and, more broadly in insect-plant interaction in nocturnal pollinators.

## Ethics

This work did not require ethical approval from humans or animal welfare committee.

## Data accessibility

Our code is publicly available without restriction, other than citation, on Github at https://github.com/MeghnaAnand5/PiliferEphysAnalysis and https://github.com/TanviDeora/PiliferBehavior. Data will be made available at figshare.

## Declaration of AI use

AI tools were used to assist with code development, identify appropriate software packages for data analysis, and edit the manuscript to meet the required word count.

## Author contributions

TD, BWB & TLD: conceptualization and design of the study; MT, AIW, TD, & TLD: data collection; MA, TD, AIW & TLD: data analysis, MA, TD, AIW, BWB & TLD wrote the manuscript, TD, BWD & TLD provided financial support.

## Acknowledgments

We thank Eatai Roth for sharing the videos of hawkmoths tracking flowers and help analyzing the data and Simon Sponberg for discussions during the design of the study. We also thank the Core Imaging Facility (CIF) of the Department of Life Sciences, SNS, managed by the Shiv Nadar IoE DTU Delhi-NCR and funded by the DST-FIST (Grant No. SR/FST/LS-1/2017/59(c)) for the use of the confocal microscopes. We thank Wai Pang Chan and the electron microscopy facility in the Department of Biology, at the University of Washington for access to the JCM-6000Plus (JEOL, Japan) scanning electron microscope.

