## Supplementary Figures and Methods for "Pilifers provide rapid mechanosensory feedback during floral tracking in hawkmoths"

### Supplementary Information for Pilifers provide rapid mechanosensory feedback during floral tracking in hawkmoths.

#### 1 Supplementary Figures

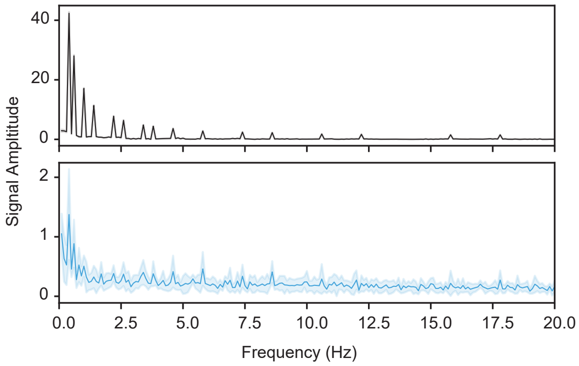


Figure 1: **Frequency peaks in floral movement and probsocis bending.** Fast Fourier Transform of the flower motion (in black) and proboscis deflection (in blue) showing the frequencies at which the flower is moving and the consequent frequencies present in the proboscis deflection.

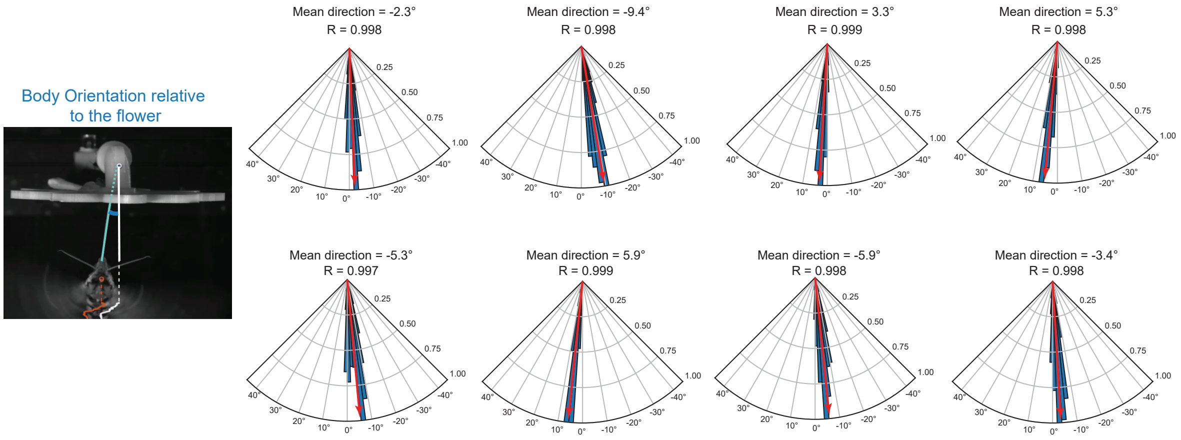


Figure 2: **Orientation of the moth with respect to the moving flower.** Although for each flight bout, the hawkmoth body approaches the flower at a slightly different angle, they tightly maintain this orientation during the entire flight bout (vector strength of mean orientation is >0.99 with a p values of $< 0.05$ across all 8 animals, Rayleigh's test). The illustration on the left shows how we computed the orientation angle (blue) relative to the flower. Each rose plot shows the distribution of orientation during a single flight bout for an individual animal (blue). The direction of the red arrow represents the mean orientation and it's length (R) represents the strength of the mean orientation. A vector strength of 1 suggests that the orientation was perfectly maintained, smaller values suggest a greater variation.

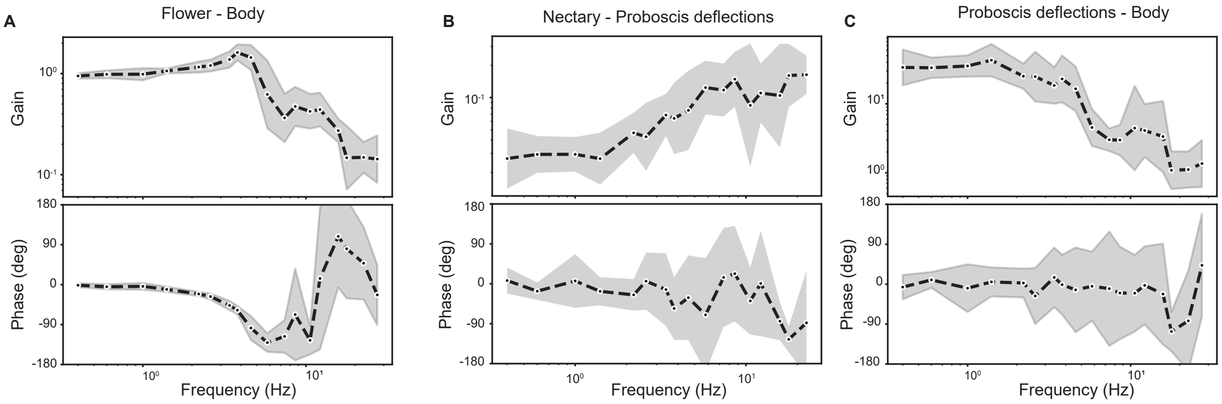


Figure 3: **Bode plots of floral motion, proboscis bending and body movements.** Bode plots showing the gain and phase between the A) Flower and body movements, B) flower and deflections at the proboscis base, and C) proboscis deflection and body movements.

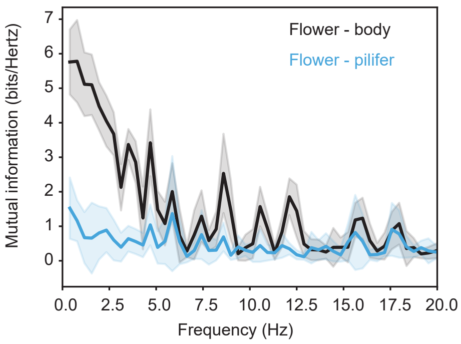


Figure 4: **Mutual information between flower, proboscis bending and body movements.**

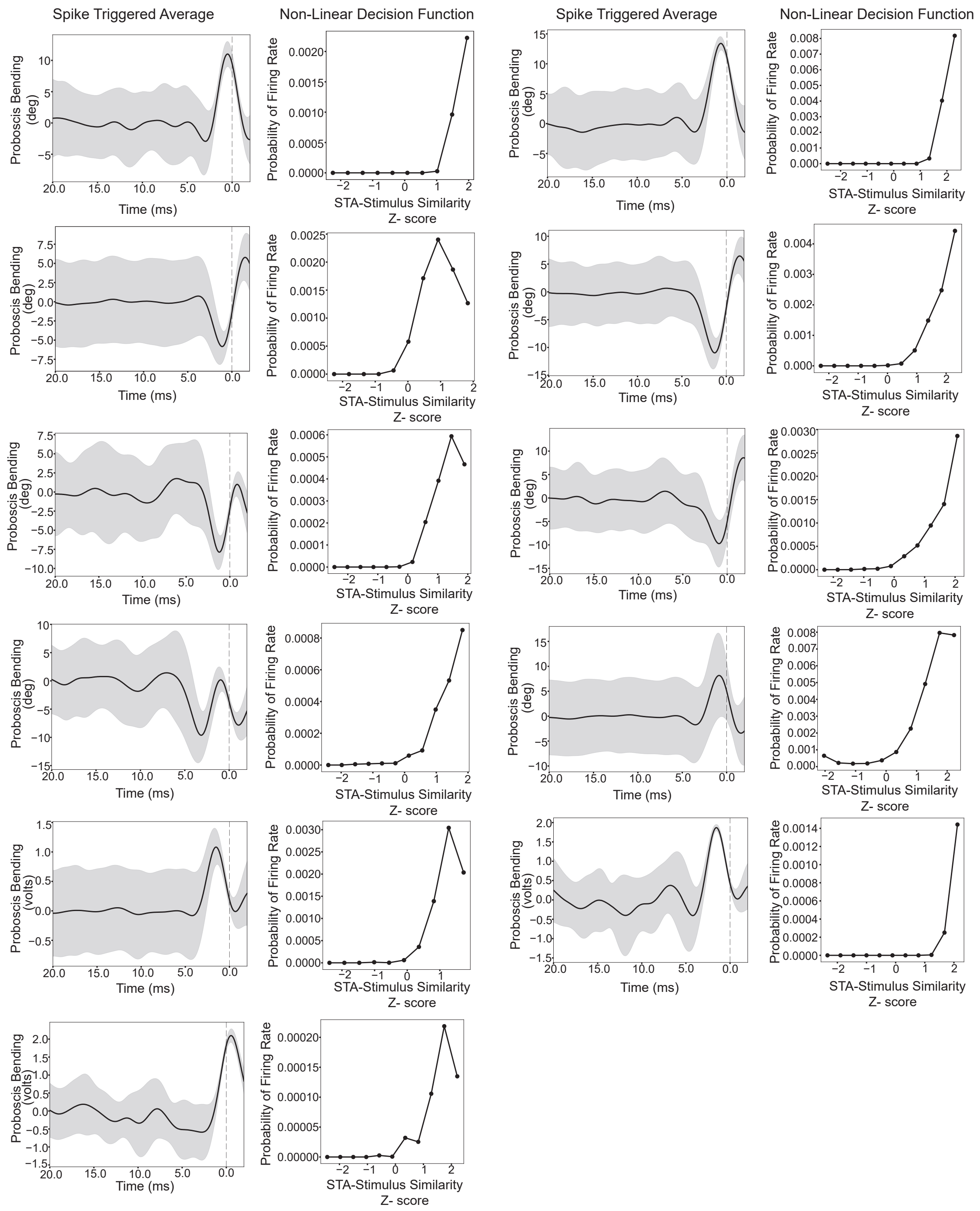


Figure 5: The spike triggered average (STA, on the left) and non linear decision function (NLD, on the right)for each individual unit.

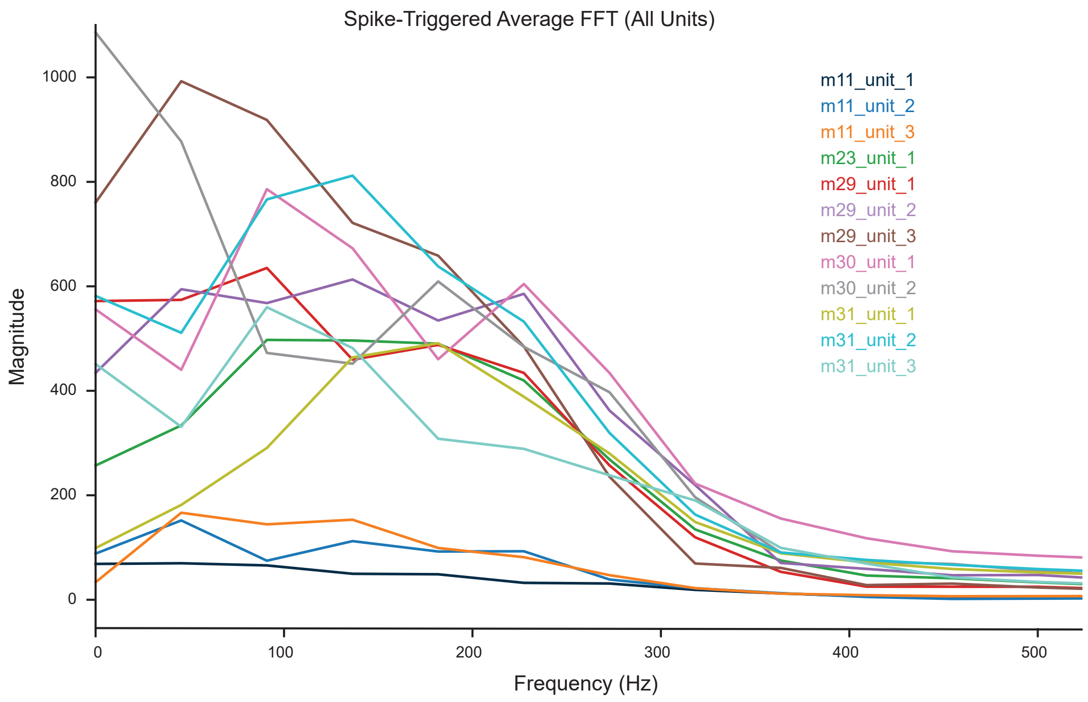


Figure 6: **Frequency tuning of individual units.** Each color represents an individual unit. The plot shows the spike triggered average in the frequency domain.

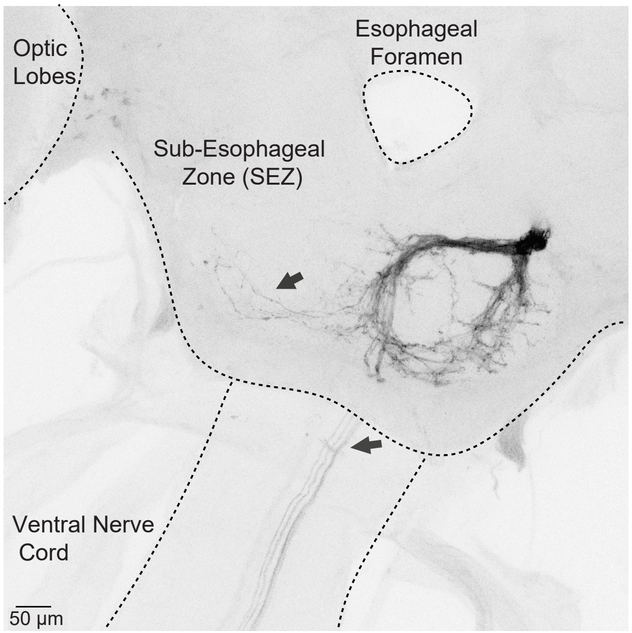


Figure 7: **Axonal projections of pilifer hair bristles.** Ipsilateral and contralateral projections into the sub-esophageal zone (SEZ) with arborizations into the ventral nerve cord (VNC) for a *Hippotion* sp.

### 2 Supplementary Methods

#### Moth Culture

Tobacco hawkmoths (*Manduca sexta*) were obtained from a colony at the University of Washington that was maintained on a 12:12 hr light:dark cycle at 20-25°C. *Hippotion sp.* were wild caught from the campus of Shiv Nadar Institution of Eminence and kept at room temperature of 25-26°C. *Daphnis nerii* eggs were obtained from a colony at the National Center for Biological Sciences. They were bred and maintained on a 12:12 hr light:dark cycle at Shiv Nadar Institution of Eminence and kept at room temperature of 25-26°C.

#### 2.1 Scanning Electron Microscopy

We fixed the hawkmoth head in 4% paraformaldehyde for 12-14 hours, and serially dehydrated the tissue by incubating it in increasing concentration of ethanol for 10 min each. We rinsed the samples twice in 100% ethanol and then critical point dehydrated the samples using Denton DCP-1 critical point dryer in the Biology Imaging Center at the University of Washington. We mounted and sputter coated whole moth heads and imaged them using a JCM-6000Plus (JEOL, Japan) SEM microscope.

#### Electrophysiology experimental setup and analysis

We used cold anesthetised *Manduca sexta* at 4°C and immobilised them in a custom made holder kept at 4°C with a stepper motor attached approximately at the proboscis knee joint. Motor commands for lateral motion to the proboscis were initiated by MATLAB via a custom built motor controller and sent to the stepper motor. A position readout signal from the motor controller was recorded by an Intan Technology RHD2000 recording system. A 16-channel silicon probe (Cambridge Neurotech P-1) was inserted at the base of the pilifer hair to record from the afferents. The probe was connected to an RHD2132 Intan Technology headstage and data was recorded at 30kS/second by the Intan technology RHD2000 recording system. We spike sorted the neural activity using the Plexon Offline Sorter across 15 animals using multiple channels within each animal. 6 animals had at least one channel with sufficient signal to noise to allow clean sorting. We set a threshold to detect firing events in either a single channel or stereodes, and used the Perform Automatic Sorting function with Valley Seeking to visualize the clusters. We further modified the clusters manually to ensure complete separation. This resulted in 16 clearly segregated units across 6 animals which were used for further analysis. We plotted a raster plot and peristimulus time histogram for each unit across the 6 animals, which showed spiking activity across 10s of stimulus for all 30 repeats. For each animal, we recorded the distance between the motor attachment and the proboscis base. We computed the lateral distance moved by the proboscis at the point it was connected to the stepper motor shaft across the voltage range used in our mechanical stimulus. In parallel, for each moth we noted the distance between the proboscis base and the point at which it was connected to the shaft. We used these two values to compute the mechanical stimulus in proboscis bending angle in degrees for each moth. We conducted a reverse correlation analysis to compute the average stimulus that led to a response. In brief, we built a spike-triggered ensemble (STE) by selecting the stimulus history 20ms preceeding each spike time. We then computed the mean of STE, which gives the spike-triggered average (STA). We also generated a prior stimulus ensemble (PSE) by selecting random timepoints across the stimulus window and selecting 20ms preceding those random timespoints. To build the nonlinear decision function (NLD), we computed a histogram of the projections of the STA onto each STE, with a bin width of 0.3 standard deviation of the PSE. Similarly, we constructed a histogram of the projections of the prior average onto each element of the PSE. We divided the normalized histogram built from the STE by the normalized histogram built from the PSE, giving us the probability of firing as a function of the similarity between the stimulus & STA.

#### Visualizing pilifer sensory projections

We performed backfills on male and female hawkmoths that were 1-5 days post-eclosion. Preference was given to healthy, active and alert moths to ensure they survived the incubation period. The moths were cold anesthetized at 4°C for 10-15 minutes, then descaled on their head, neck and thorax with a moist tissue. Moths were anesthetized again if they became active during descaling. We secured the moths in an inverted centrifuge tube using dental wax melted by an electric cauterizer. The melted wax was gently placed between the edge of the holder and both sides of the thorax, then left to solidify. The moth was thereby held in place with its head protruding from the holder. We used a Nikon SMZ800N microscope at about 8X magnification to perform backfills. The syringe tip from a 1ml insulin syringe was used to uniformly shave all pilifer hairs visible on the cuticle, on either side at the base of the proboscis. Immediately after, we placed a drop of distilled water on the shaved cuticle using the same syringe tip. Dextran Texas Red 3000 MW (Invitrogen by Thermo Fisher Scientific, lysine fixable, D3328) crystals were minimally diluted with distilled water and added to the pilifer cuticle in quick succession. We waited for 30-60 seconds for absorption and repeated the procedure 3-5 times. Moth were fed 20% concentrated sugar solution and placed in a moist chamber and incubated for 24-72 hours at 25-30°C. After incubation, the moth was anesthetized at -20°C for 15 minutes and the head and abdomen were placed in 4% paraformaldehyde and fixed for 12-14 hours depending on the size of the moth. We dissected the brain and ventral nerve cord under a Nikon SMZ745N microscope. We serially dehydrated the tissue by incubating it in 50%, 60%, 75%, 90% ethanol and twice in 100% ethanol for 10 mins sequentially. After the final ethanol wash we added 2ml of methylsalicylate until complete clearing of the sample. We then mounted it on a microscope slide in methylsalicylate for confocal laser microscopy imaging. We imaged it with either the Nikon Ti2E, A1R MP or Nikon Ti2E, AXR-NSPARC confocal microscopes. The images were obtained at 10X, 20X or 40X. Images were analyzed in Fiji software.
